# Distinct Patterns of the Autofluorescence of Body Surface: Potential Novel Diagnostic Biomarkers for Stable Coronary Artery Disease and Myocardial Infarction

**DOI:** 10.1101/330985

**Authors:** Xinkai Qu, Yujia Li, Yue Tao, Mingchao Zhang, Danhong Wu, Shaofeng Guan, Wenzheng Han, Weihai Ying

**Affiliations:** Department of Cardiology, Huadong Hospital Affiliated to Fudan University, Shanghai 200040, P.R. China; Med-X Research Institute and School of Biomedical Engineering, Shanghai Jiao Tong University, Shanghai 200030, P.R. China; Department of Cardiology, Shanghai Chest Hospital Affiliated to Shanghai Jiao Tong University, Shanghai 200030, P.R. China; Department of Neurology, Shanghai Fifth People’s Hospital Affiliated to Fudan University, Shanghai 200240, P.R. China; Collaborative Innovation Center for Genetics and Development, Shanghai 200043, P.R. China

**Keywords:** Autofluorescence, Coronary artery disease, Myocardial infarction.

## Abstract

Searches for new biomarkers of stable coronary artery disease (SCAD) and myocardial infarction (MI) are critical for therapeutic efficacy of the diseases. In this study we tested our hypothesis that distinct patterns of autofluorescence (AF) of skin and fingernails may become novel diagnostic biomarkers for MI and SCAD. Our study has indicated that SCAD and MI have distinct patterns of AF of their body surface: First, the AF intensity of the MI patients is significantly higher than that of the Healthy and Low-Risk group in their right and left Centremetacarpus, Ventroforefinger, Dorsal Index Finger and Ventribrachium, while the AF intensity of the SCAD patients is significantly higher than that of the Healthy and Low-Risk group in their right and left Index Fingernails and Dorsal Antebrachium; and second, the AF asymmetry of the MI patients is significantly higher than that of the Healthy and Low-Risk group in their Centremetacarpus, Ventroforefinger, Index Fingernails and Dorsal Antebrachium, while the AF asymmetry of the SCAD patients is significantly higher than that of the Healthy and Low-Risk group in their Ventroforefinger, Dorsal Index Finger, Dorsal Centremetacarpus and Index Fingernails. Moreover, the AF pattern of acute ischemic stroke is markedly different from those of SCAD and MI. The oxidative stress in the plasma of the MI and SCAD patients may cause the increased AF by altering the AF of keratins. Collectively, our study has indicated that SCAD and MI patients have distinct patterns of AF changes, which may become novel diagnostic biomarkers for SCAD and MI.

## Introduction

Cardiovascular diseases remain the leading cause of mortality globally with an estimated 30% of adults living with the diseases. It is projected that more than 23.3 million people around the world will die annually from cardiovascular diseases including acute myocardial infarction (AMI) and stroke by 2030 (10). Stable coronary artery disease (SCAD) refers to patients with known obstructive or nonobstructive coronary artery disease whose symptoms are stable on medical therapy. Clinical risk factors and risk scores are frequently used to predict cardiovascular events in SCAD patients, which often lack sensitivity and specificity (8).

Acute coronary syndrome (ACS) describes the myocardial ischemic states including unstable angina (UA), ST-elevated myocardial infarction (STEMI) and non-ST elevated myocardial infarction (NSTEMI). The diagnosis and classification of ACS is based on clinical assessment, electrocardiogram (ECG) findings and biochemical markers of myocardial necrosis including increased levels of cardiac troponins (16,18). The term of ‘myocardial infarction’ (MI) is used when there is myocardial necrosis in the setting of acute myocardial ischemia. The presence of persistent ECG findings of ST segment elevation is used to differentiate STEMI from NSTEMI (15). MI may cause heart failure, an irregular heartbeat, cardiogenic shock, or cardiac arrest. Approximately 15.9 million MI occurred in 2015 globally (21). More than 3 million people had an STEMI, while more than 4 million people had an NSTEMI. The death rate for an STEMI patients in the developed world is approximately 10% (17).

Due to the severity and high incidence of MI and SCAD, it is necessary to search for the novel approaches that can diagnose MI and SCAD non-invasively, efficiently and economically. Early diagnosis of AMI is of great clinical significance: Delayed confirmation of the diagnosis of an AMI may increase the AMI-associated complications (19),while a delay in ruling out the diagnosis can lead to overcrowding in the emergency department (16). An ideal approach should enable patients to diagnose MI and SCAD at home, which can not only greatly decrease the burden on hospitals, but also greatly enhance the probability of early diagnosis of the diseases. However, so far the only potential approach that may be used for home-based diagnosis of MI is ECG, if ECG can be supported by big data and artificial intelligence (AI) technology. However, ECG is often insufficient to diagnose AMI, since ST-segment deviation may be observed in other pathological conditions such as left ventricular hypertrophy, acute pericarditis and left bundle-branch block (16).

Human autofluorescence (AF) has been used for non-invasive diagnosis of diabetes, which is based on detection of the AF of advanced glycation end-products (AGEs) of the collagen in dermis (11,14). It is of significance to further investigate the potential of AF as biomarkers for diseases, due to the non-invasiveness and simplicity of AF detection. Our recent study has suggested that UV-induced epidermal green AF, which is generated from UV-induced keratin 1 proteolysis in the spinous cells of epidermis, is the first non-invasive biomarker for predicting UV-induced skin damage (7). We have further found that the oxidative stress induced by UVC causes the increased epidermal AF by inducing keratin 1 proteolysis (12). Since there are significant increases in oxidative stress in the plasma, serum or urine of SCAD patients (1,2,5,6,20) and MI patients (3,4,9,13,20), it is warranted to determine if there are increases in the epidermal AF of SCAD and MI patients, which may become novel diagnostic biomarkers for the diseases.

In this study, we tested our hypothesis that SCAD and MI patients may have increased AF in their fingernails and certain regions of their skin. Our study has provided first evidence that SCAD and MI patients have distinct patterns of green AF in their fingernails and certain regions of the skin, suggesting that the distinct AF patterns may become new diagnostic biomarkers for SCAD and MI.

## Methods

### Human subjects

The study was conducted according to protocols approved by the Ethics Committee of Shanghai Fifth People’s Hospital Affiliated to Fudan University, and by the Ethics Committee of Shanghai Chest Hospital Affiliated to Shanghai Jiao Tong University. The human subjects in our study were divided into four groups. Group 1: The group of healthy persons and Low-Risk persons - the persons with only a mild level of hypertension; Group 2: The group of SCAD patients – the people who were clinically diagnosed as SCAD patients by the cardiologists of the Department of Cardiology, Shanghai Chest Hospital Affiliated to Shanghai Jiao Tong University; Group 3: The group of MI patients – the people who were clinically diagnosed as MI patients by the cardiologists of the Department of Cardiology, in which 5 of the 8 MI patients were AMI patients, while 3 of the 8 MI patients were stable phase post-MI patients, and Group 4: The group of acute ischemic stroke (AIS) patients who had ischemic stroke within 7 days from the time of examination, who were clinically diagnosed as AIS patients by the neurologists of the Department of Neurology, Shanghai Fifth People’s Hospital Affiliated to Fudan University. The age of Group 1, Group 2, Group 3 and Group 4 is 67.90 ± 7.92, 65.40 ± 9.25, 62.38 ± 15.15 and 67.30 ± 10.27 years of old, respectively.

### Autofluorescence determination

For all of the human subjects, the AF intensity in the following seven regions on both hands, i.e., fourteen regions in total, was determined, including the index fingernails, Ventroforefingers, dorsal index fingers, Centremetacarpus, Dorsal Centremetacarpus Ventriantebrachium, and Dorsal Antebrachium. A portable AF imaging equipment was used to detect the AF of the these fourteen regions of the human subjects. The excitation wavelength is 485 nm, and the emission wavelength is 500 - 550 nm.

### Statistical analyses

All data are presented as mean <u>±</u> SEM. Data were assessed by one-way ANOVA, followed by Student - Newman - Keuls *post hoc* test, except where noted. *P* values less than 0.05 were considered statistically significant.

## Results

We determined the green AF intensity of the Centremetacarpus of the Healthy and Low-Risk group, the SCAD group, the MI group, and the AIS group. All of these patient groups have higher AF intensity in this region than the Healthy and Low-Risk group, except that the SCAD group and the Healthy and Low-Risk group have the same AF intensity at the left Centremetacarpus (Fig. 1A). We further determined the AF asymmetry at the Centremetacarpus of the Healthy and Low-Risk group, the SCAD group, the MI group, and the AIS group. Both the MI group and AIS group have significantly higher AF asymmetry than the Healthy and Low-Risk group in the Centremetacarpus (Fig. 1B). In the Ventroforefingers, all of the patient groups have higher AF intensity than the Healthy and Low-Risk group, except that the SCAD group and the Healthy and Low-Risk group have the same AF intensity in the region (Fig. 2A). Moreover, all of the patient groups have significantly higher AF asymmetry than the Healthy and Low-Risk group in the Ventroforefingers (Fig. 2B).

**Fig. 1.**
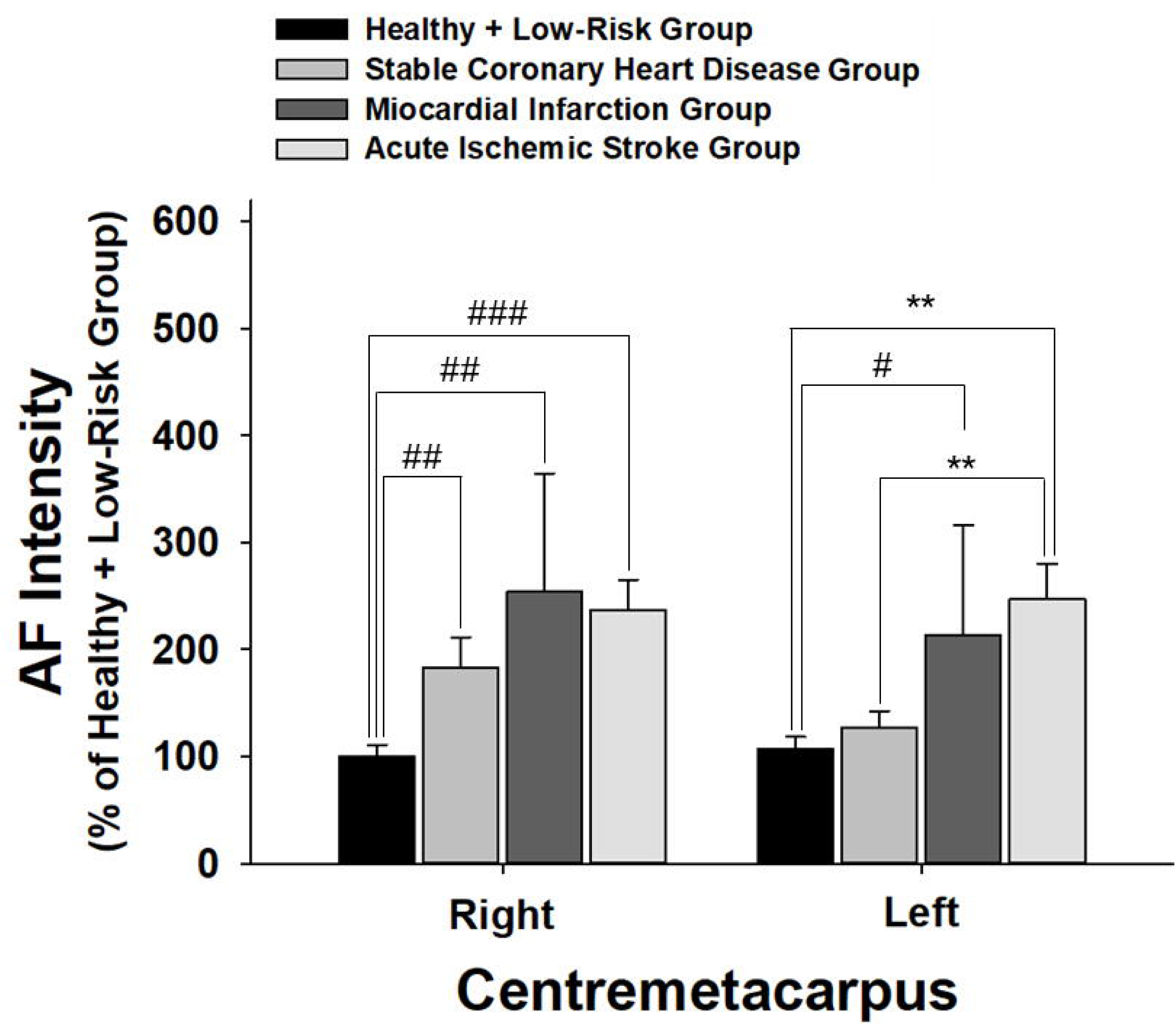
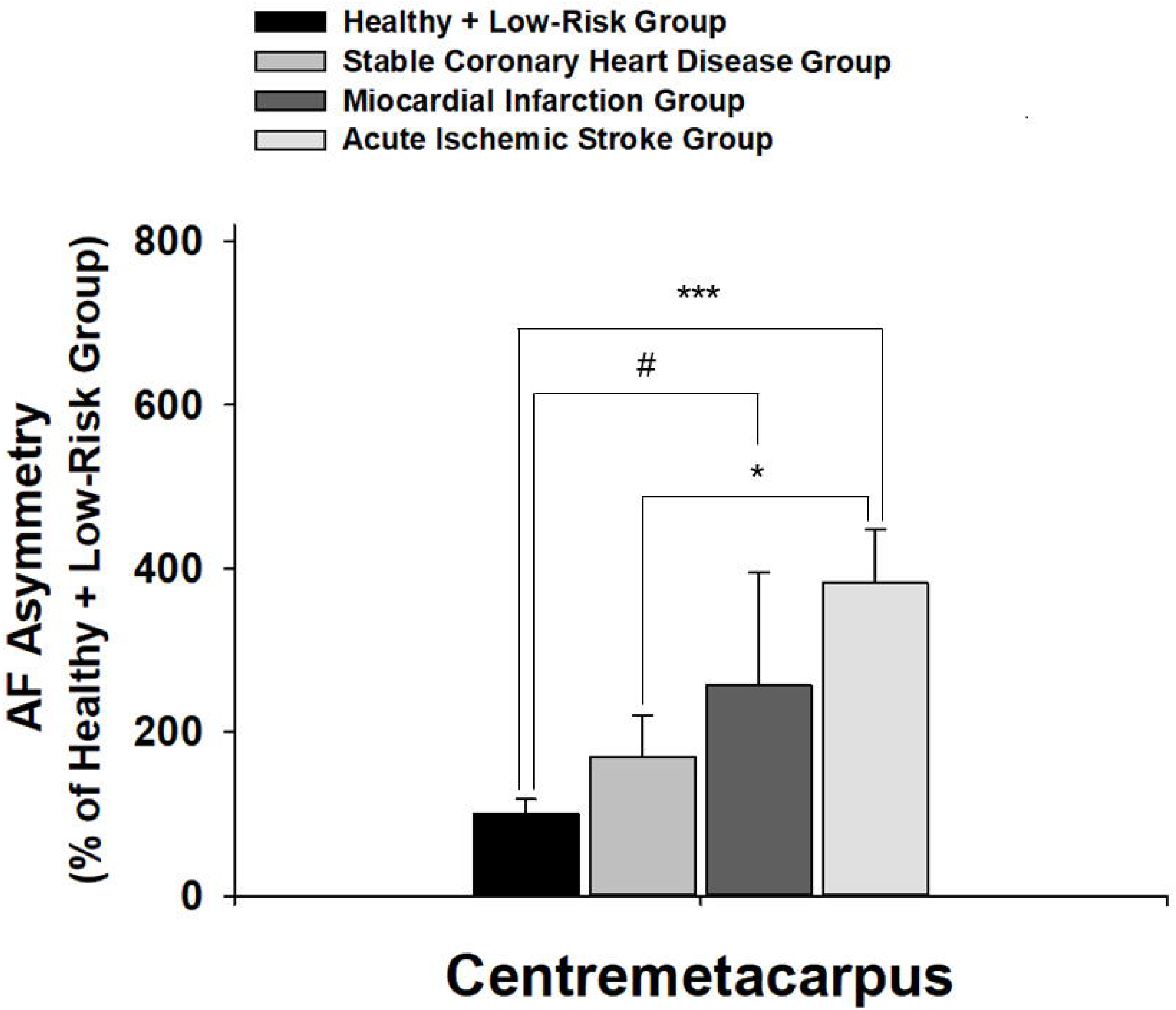
On both right and left Centremetacarpus, the MI patients and the AIS patients have significantly higher green AF intensity than the Healthy and Low-Risk persons, while the SCAD patients have significantly higher green AF intensity than the Healthy and Low-Risk persons only at right Centremetacarpus. (B) The MI patients and the AIS patients have significantly higher AF asymmetry than the Healthy and Low-Risk persons at Centremetacarpus, while there is no difference in the AF asymmetry between the SCAD group and the Healthy and Low-Risk group at Centremetacarpus. The number of subjects in the Healthy and Low-Risk group, the SCAD group, the MI group and the AIS group is 43, 20, 8, and 49-50, respectively. *, *p* < 0.05; ***, *p* < 0.001; #, *p* < 0.05 (Student *t-t*est).

**Fig. 2.**
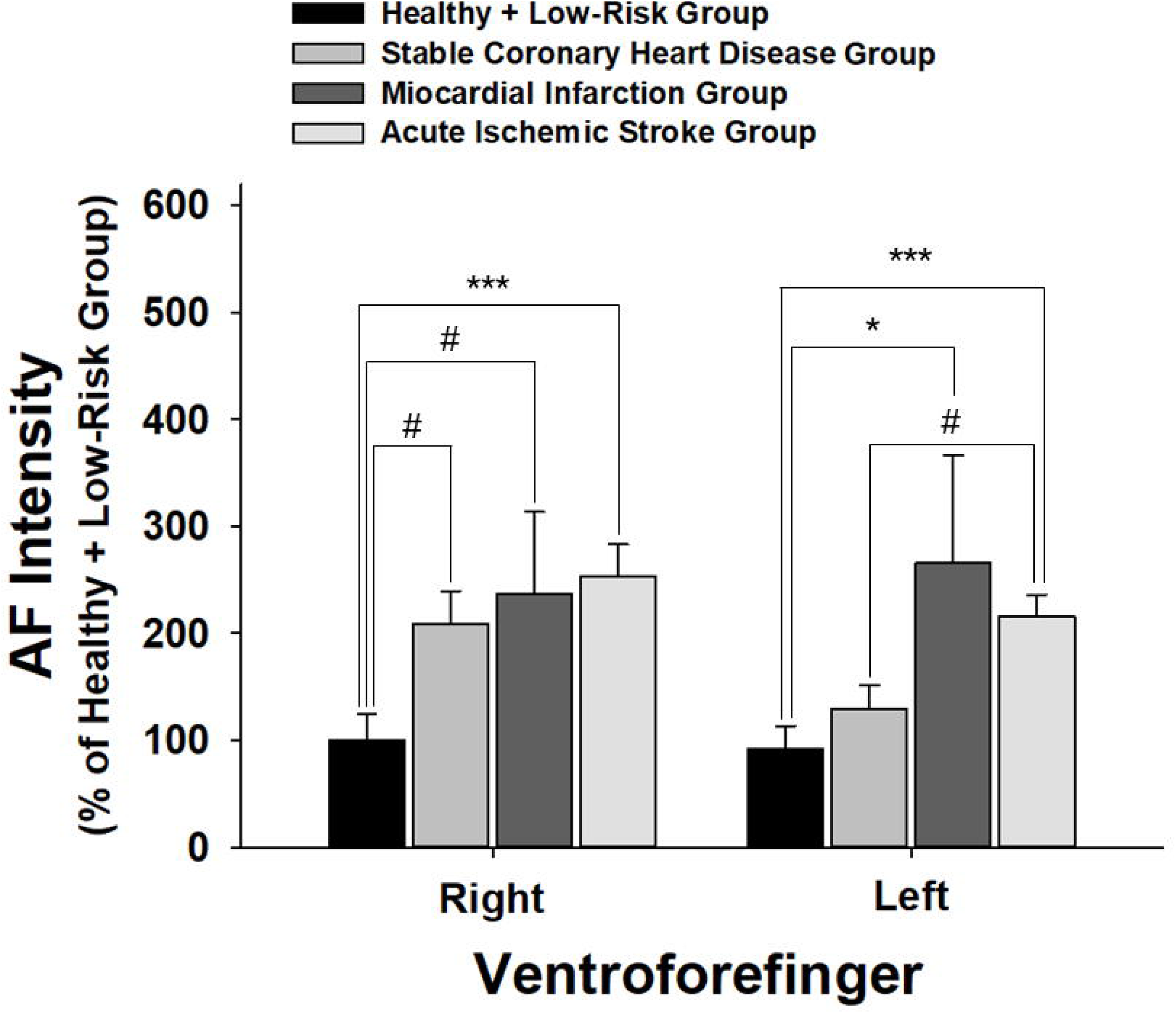
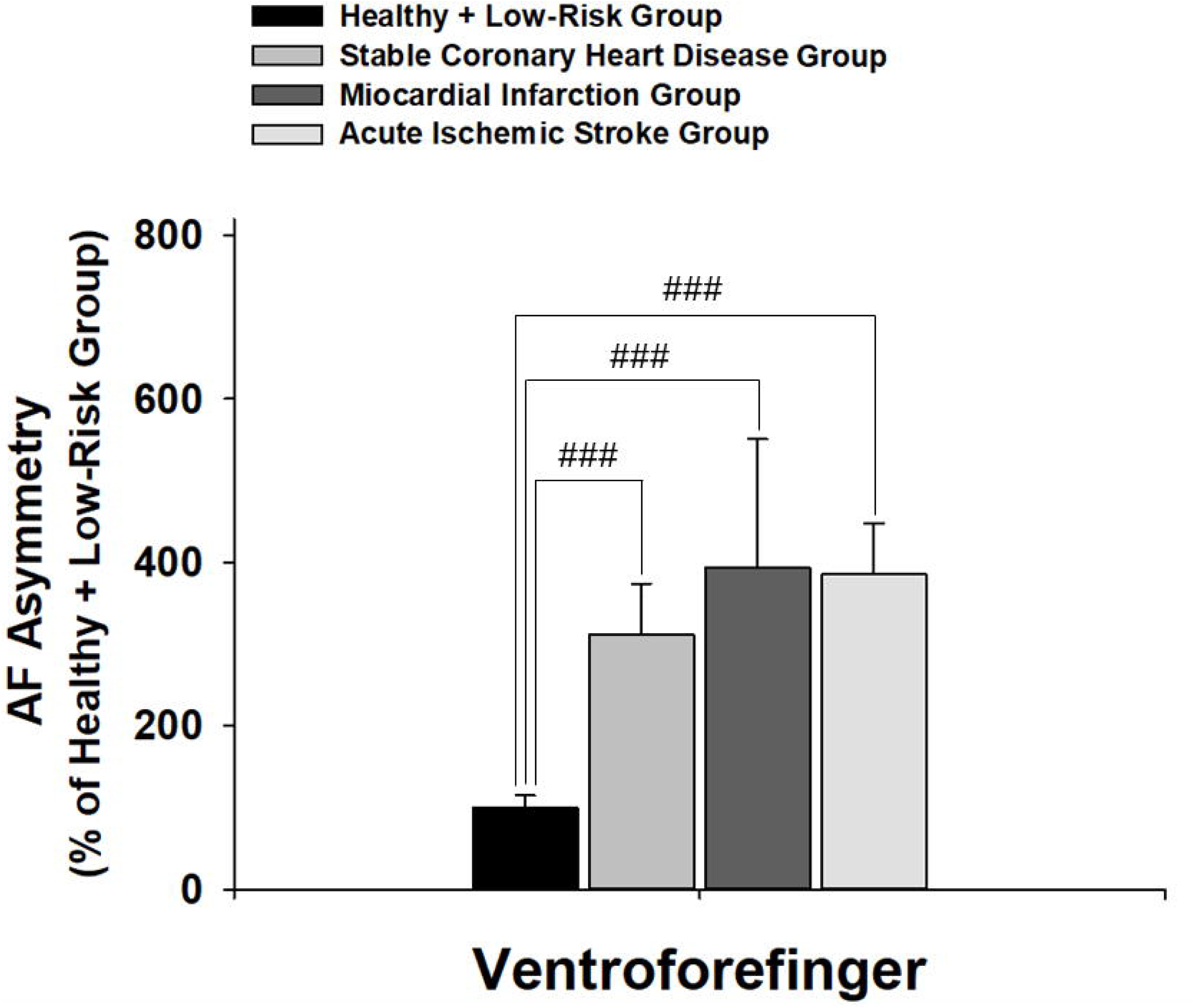
(A) On right and left Ventroforefingers, both the MI patients and the AIS patients have significantly higher green AF intensity than the Healthy and Low-Risk persons, while the SCAD patients have significantly higher green AF intensity than the Healthy and Low-Risk persons only at right Ventroforefinger. (B) The SCAD patients, the MI patients and the AIS patients have significantly higher AF asymmetry than the Healthy and Low-Risk persons at Ventroforefinger. The number of subjects in the Healthy and Low-Risk group, the SCAD group, the MI group and the AIS group is 43, 20, 8, and 49-50, respectively. ###, *p* < 0.001 (Student *t*-test).

In the Dorsal Index Fingers, all of the patient groups have higher AF intensity than the Healthy and Low-Risk group, except that the SCAD group and the Healthy and Low-Risk group have the same AF intensity in the region (Fig. 3A). Both the SCAD group and the AIS group have significantly higher AF asymmetry than the Healthy and Low-Risk group in the Dorsal Index Fingers (Fig. 3B). Only in the right Dorsal Centremetacarpus, all of the patient groups have significantly higher AF intensity than the Healthy and Low-Risk group (Fig. 4A). Both the SCAD group and the AIS group have significantly higher AF asymmetry than the Healthy and Low-Risk group in Dorsal Centremetacarpus (Fig. 4B).

**Fig. 3.**
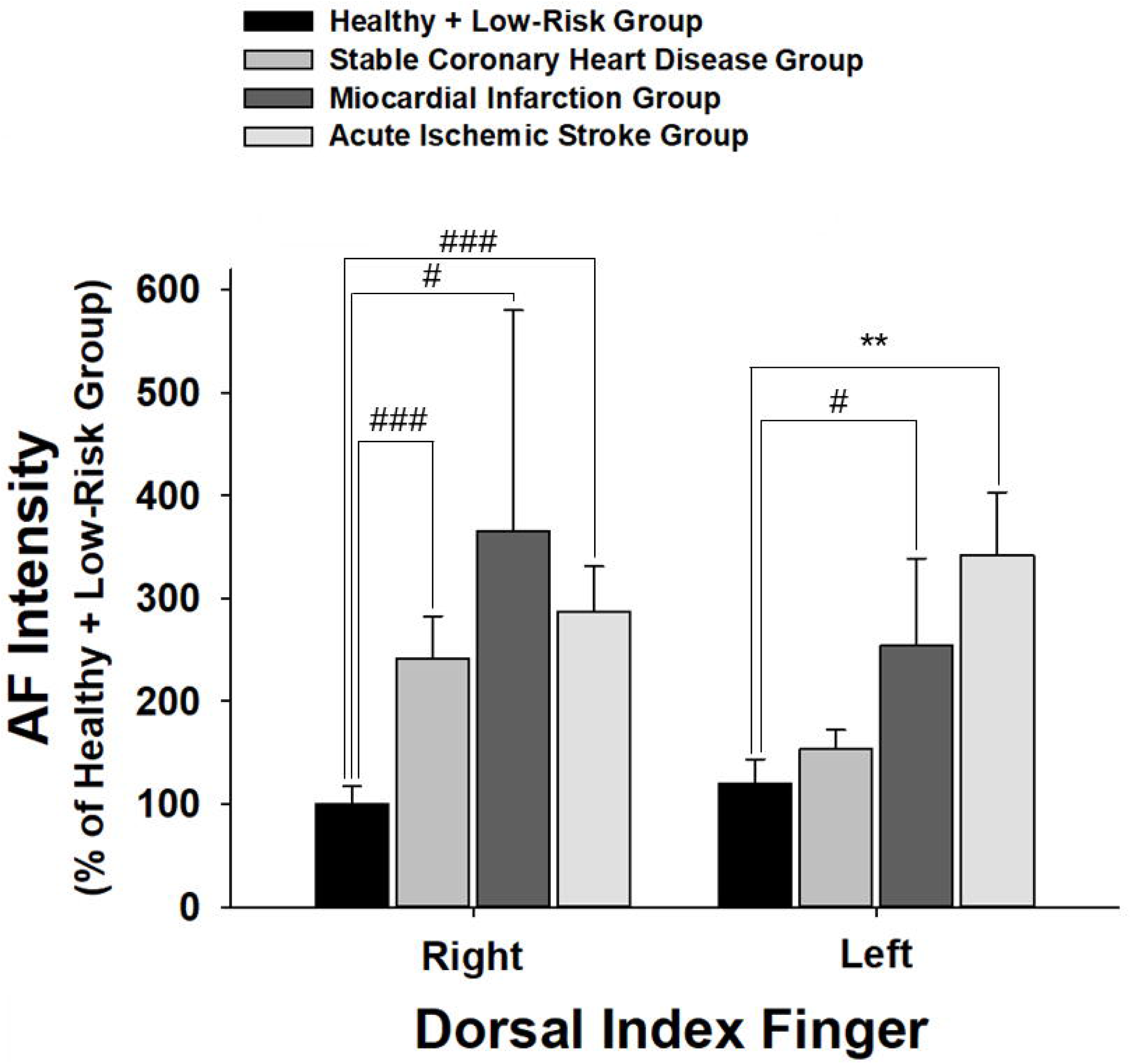
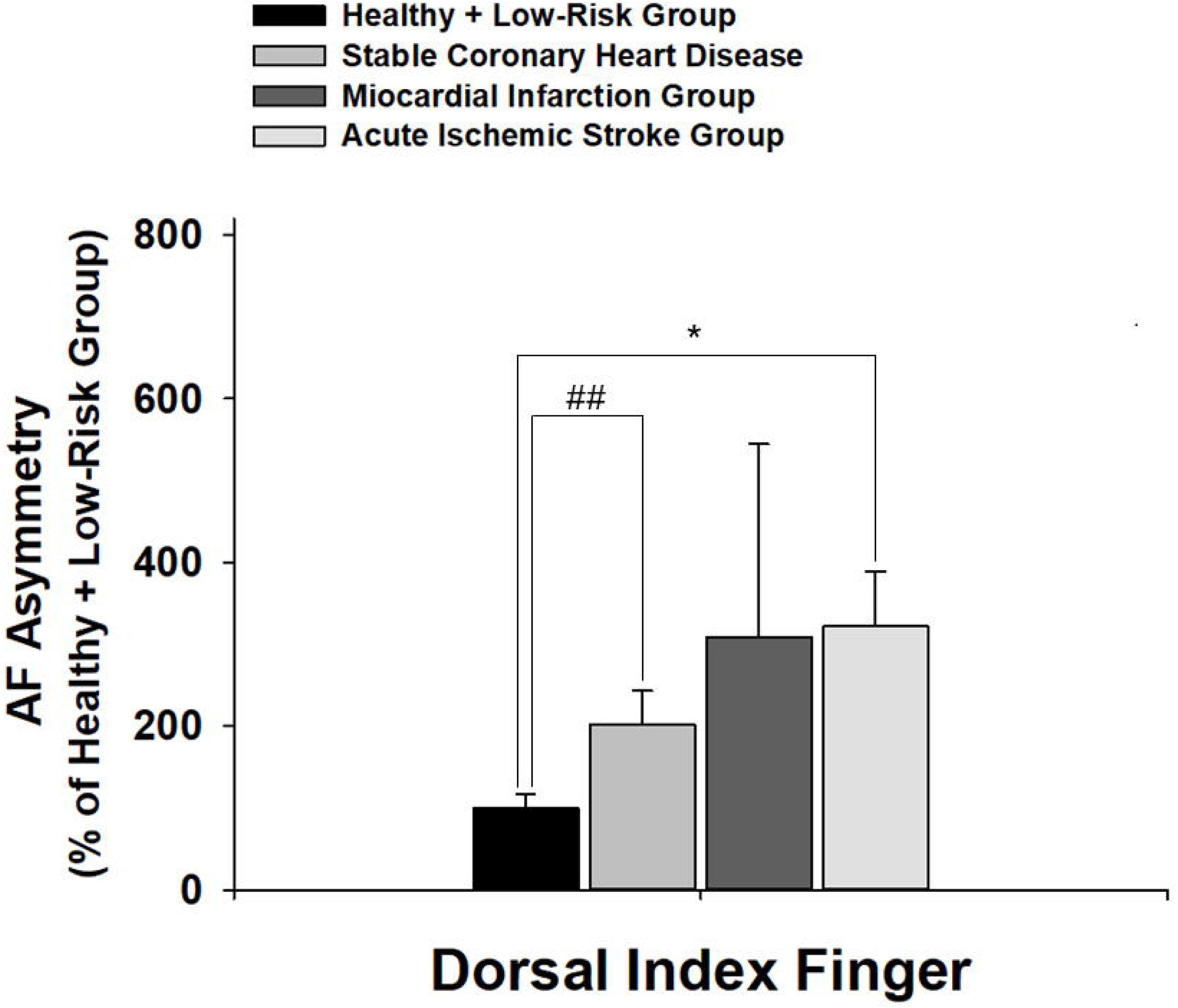
(A) On right and left Dorsal Index Fingers, both MI patients and AIS patients have significantly higher green AF intensity than Healthy and Low-Risk persons. SCAD patients have significantly higher green AF intensity than Healthy and Low-Risk persons only at right Dorsal Index Fingers. (B) The SCAD patients and the AIS patients, but not MI patients, have significantly higher AF asymmetry than the Healthy and Low-Risk persons at Dorsal Index Fingers. The number of subjects in the Healthy and Low-Risk group, the SCAD group, the MI group and the AIS group is 43, 20, 8, and 50, respectively. *, *p* < 0.05; **, *p* < 0.01; #, *p* < 0.05 (Student *t-t*est); ##, *p* < 0.01 (Student *t-t*est); ###, *p* < 0.001 (Student *t-t*est).

**Fig. 4.**
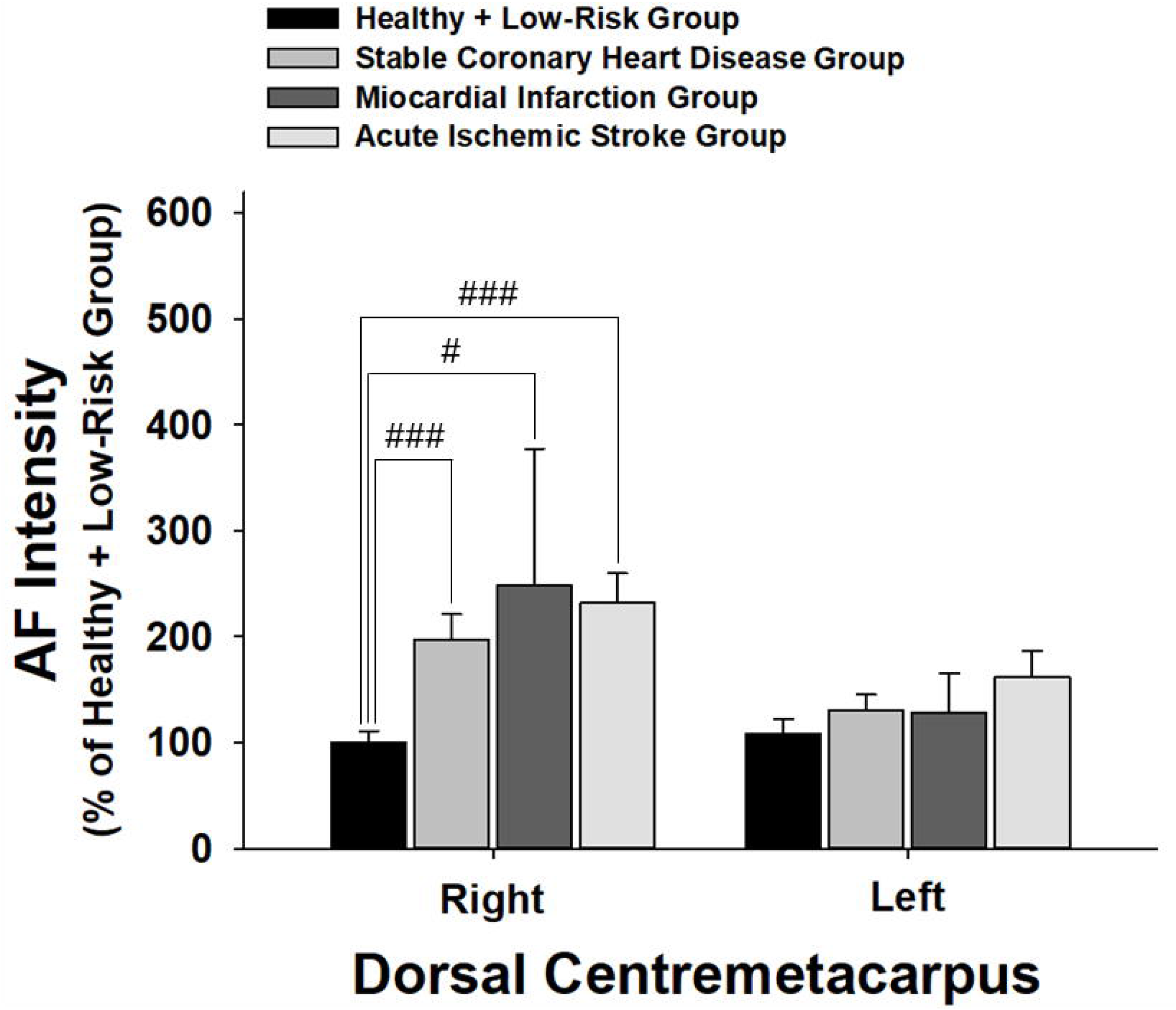
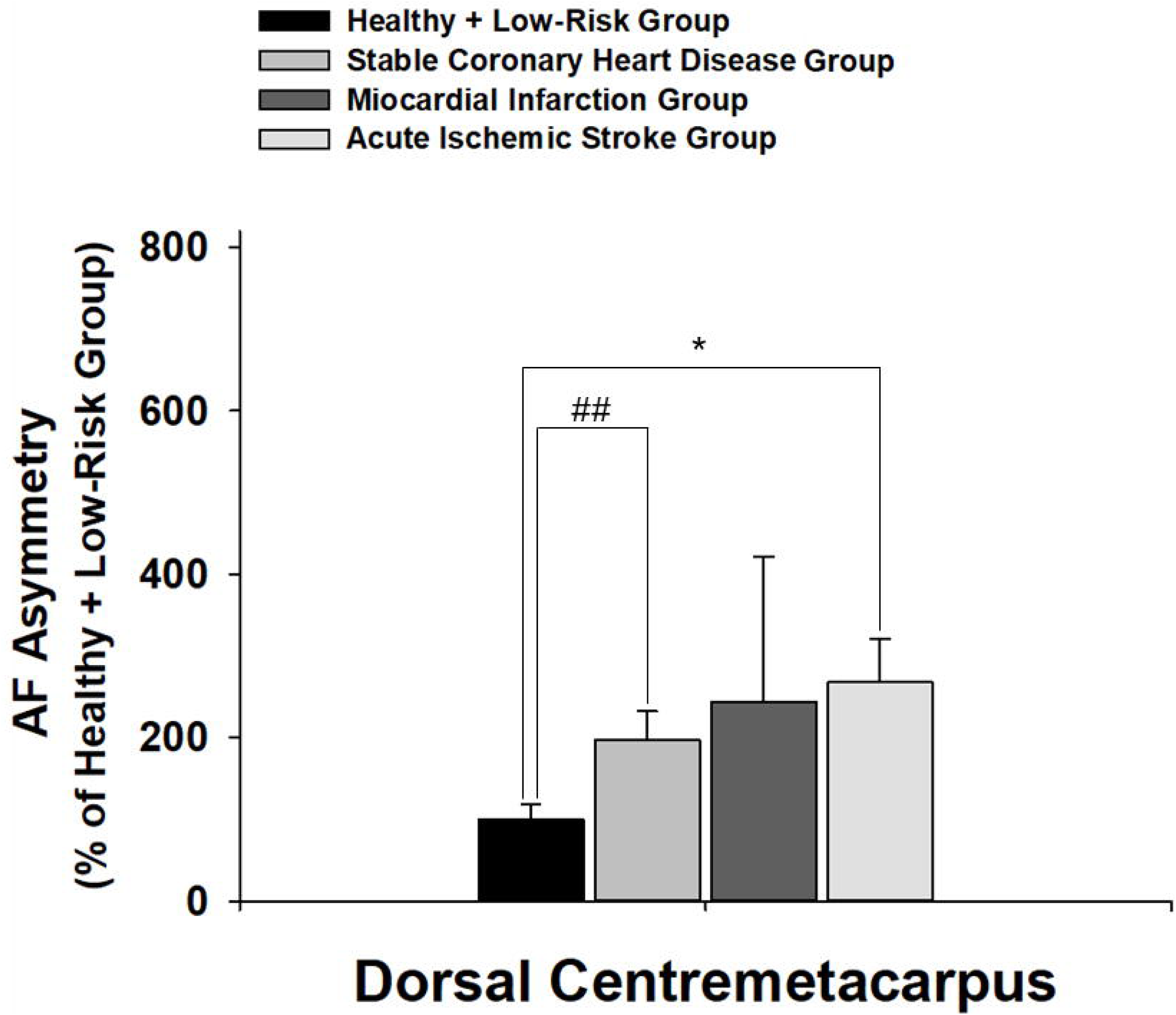
(A) Only on right Dorsal Centremetacarpus, the SCAD patients, the MI patients and the AIS patients have significantly higher green AF intensity than the Healthy and Low-Risk persons. (B) Both the SCAD patients and the AIS patients have significantly higher AF asymmetry than the Healthy and Low-Risk persons at Dorsal Centremetacarpus. The number of subjects in the Healthy and Low-Risk group, the SCAD group, the MI group and the AIS group is 43, 20, 8, and 41-46, respectively. *, *p* < 0.05; #, *p* < 0.05 (Student *t-t*est); ##, *p* < 0.01 (Student *t-t*est); ###, *p* < 0.001 (Student *t-t*est).

In the Index Fingernails, all of the patient groups have significantly higher AF intensity than the Healthy and Low-Risk group, except that the MI group and the Healthy and Low-Risk group have the same AF intensity in the region (Fig. 5A). All of the patient groups have significantly higher AF asymmetry than the Healthy and Low-Risk group in the Index Fingernails (Fig. 5B). In Dorsal Antebrachium, all of the patient groups have higher AF intensity than the Healthy and Low-Risk group, except that the MI group and the Healthy and Low-Risk group have the same AF intensity in the region (Fig. 6A). Both the MI group and the AIS group have significantly higher AF asymmetry than the Healthy and Low-Risk group in Dorsal Antebrachium (Fig. 6B). In the Ventriantebrachium, only the MI group and the AIS group have significantly higher AF intensity than the Healthy and Low-Risk group (Fig. 7A), while only the AIS group have significantly higher AF asymmetry than the Healthy and Low-Risk group (Fig. 7B).

**Fig. 5.**
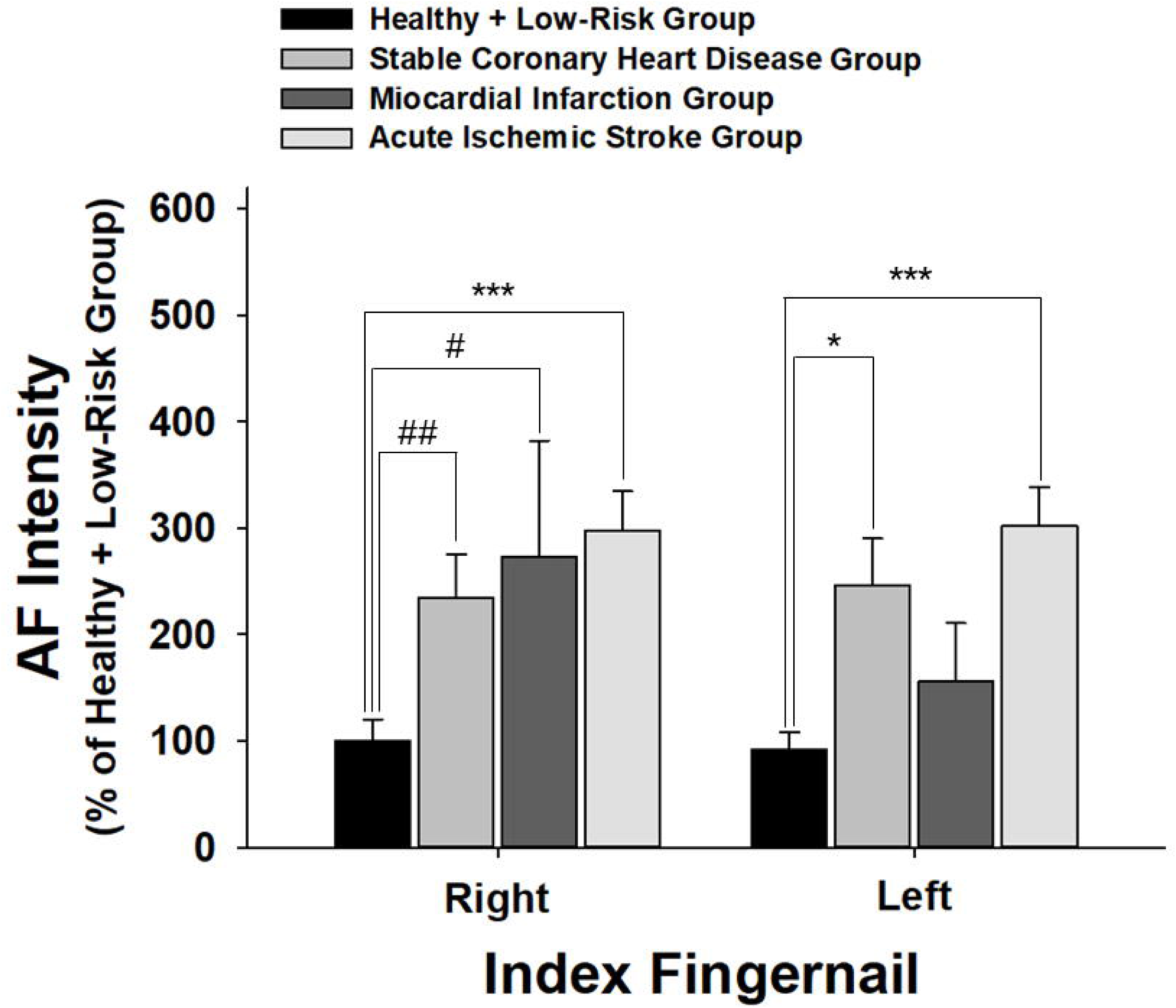
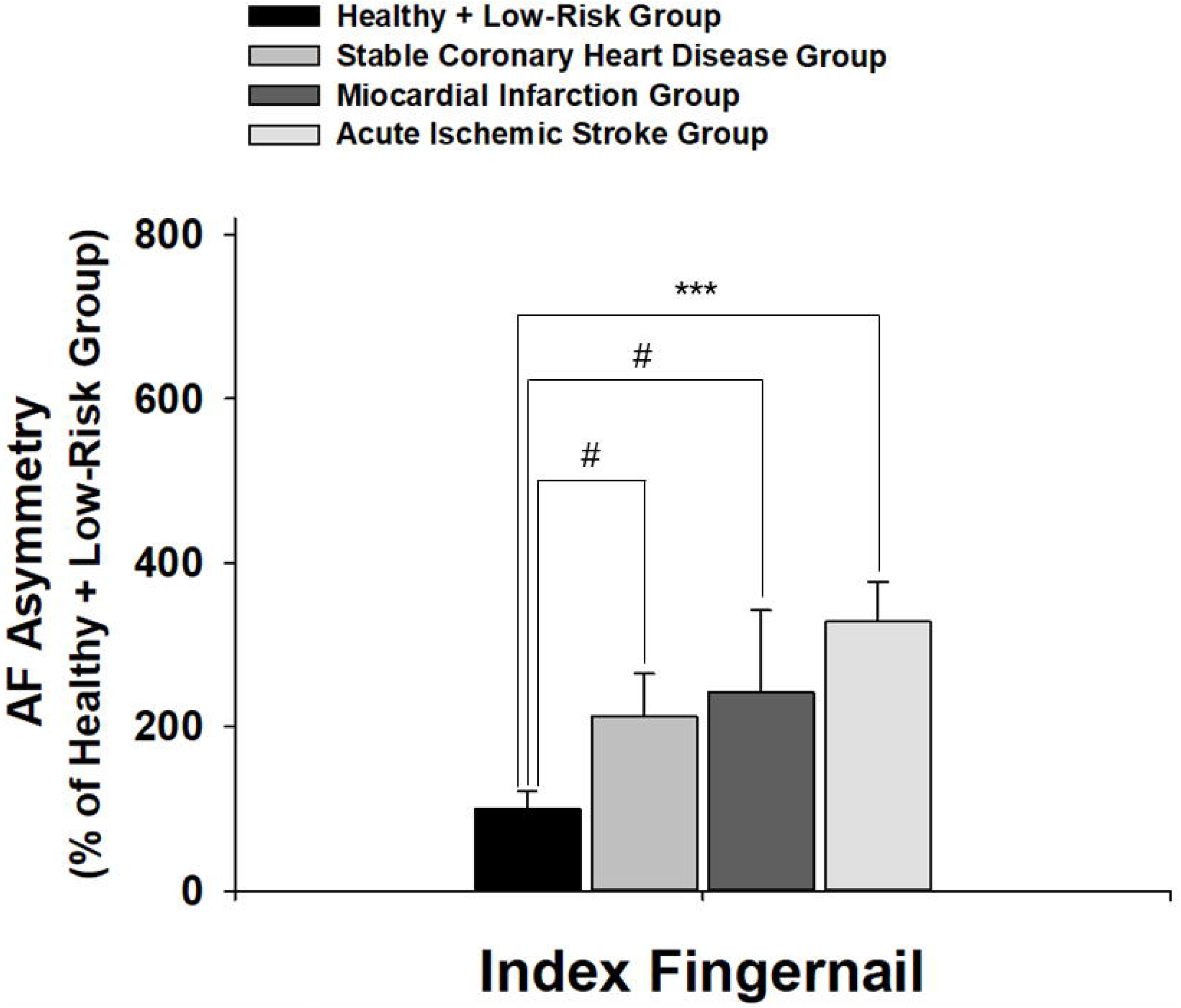
(A) On right and left Index Fingernails, both the SCAD patients and the AIS patients have significantly higher green AF intensity than the Healthy and Low-Risk persons, while the MI patients have significantly higher green AF intensity than the Healthy and Low-Risk persons only at right Index Fingernails. (B) The SCAD patients, the MI patients and the AIS patients have significantly higher AF asymmetry than Healthy and Low-Risk persons at Index Fingernails. The number of subjects in the Healthy and Low-Risk group, the SCAD group, the MI group and the AIS group is 43, 20, 8, and 50, respectively. *, *p* < 0.05; ***, *p* < 0.001; #, *p* < 0.05 (Student *t-t*est); ##, *p* < 0.01 (Student *t*-test).

**Fig. 6.**
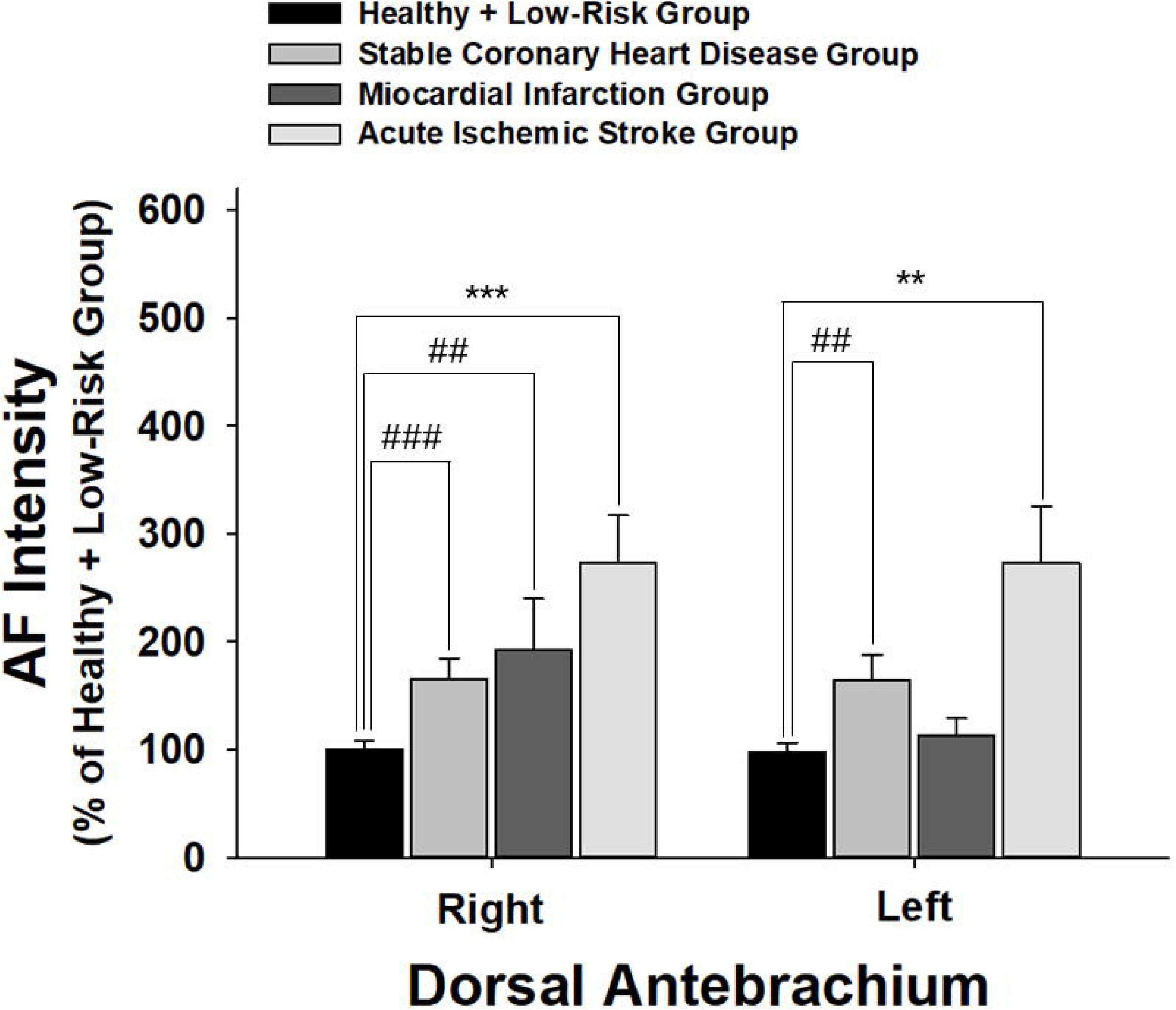
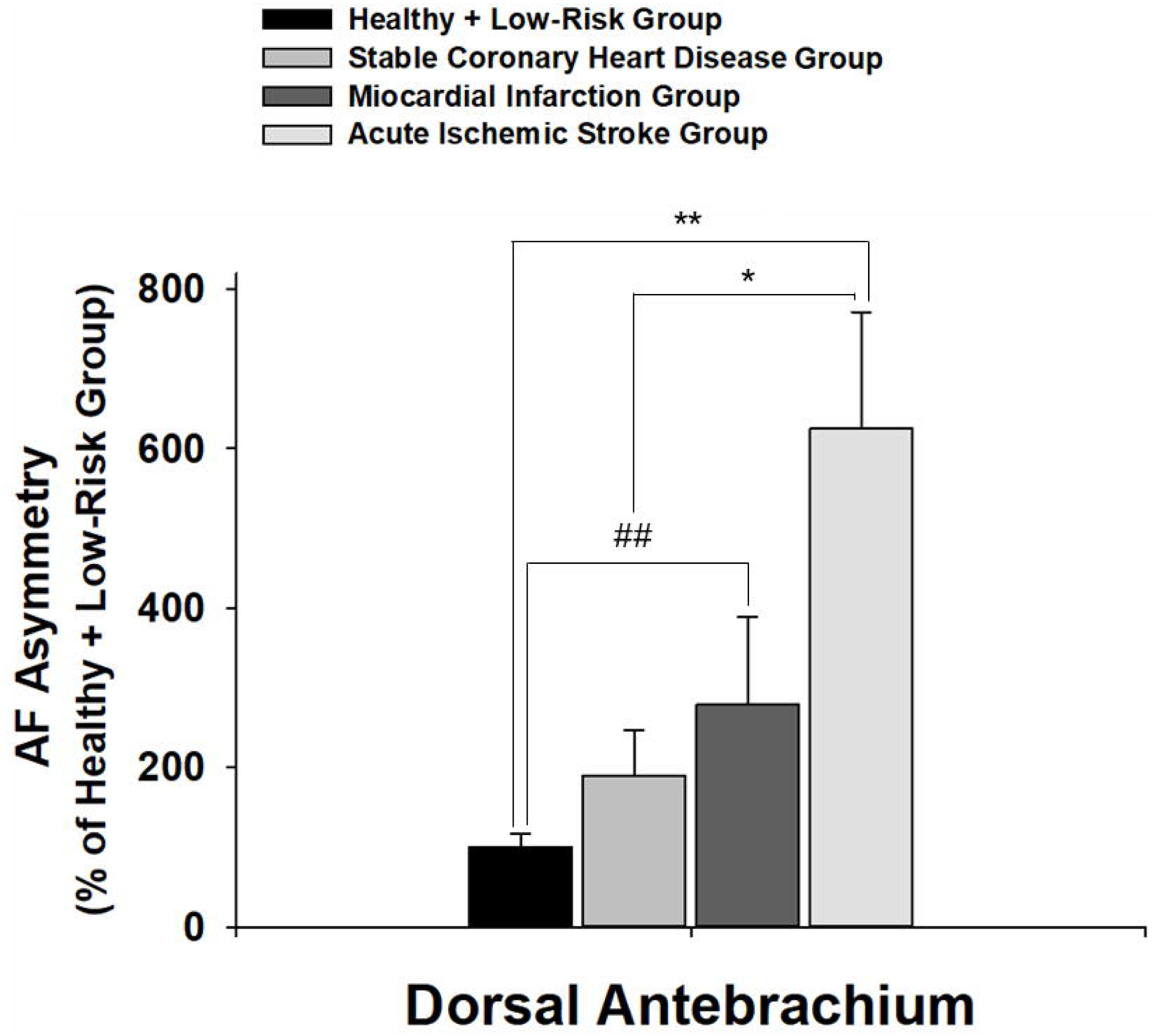
(A) On right and left Dorsal Antebrachium, both SCAD patients and AIS patients have significantly higher green AF intensity than Healthy and Low-Risk persons, while MI patients have significantly higher green AF intensity than Healthy and Low-Risk persons only at right Dorsal Antebrachium. (B) Both the MI patients and the AIS patients have significantly higher AF asymmetry than the Healthy and Low-Risk persons at Dorsal Antebrachium. The number of subjects in the Healthy and Low-Risk group, the SCAD group, the MI group and the AIS group is 43, 20, 8, and 47-48, respectively. *, *p* < 0.05; **, *p* < 0.01; ***, *p* < 0.001; ##, *p* < 0.01 (Student *t-t*est); ###, *p* < 0.001 (Student *t-t*est).

**Fig. 7.**
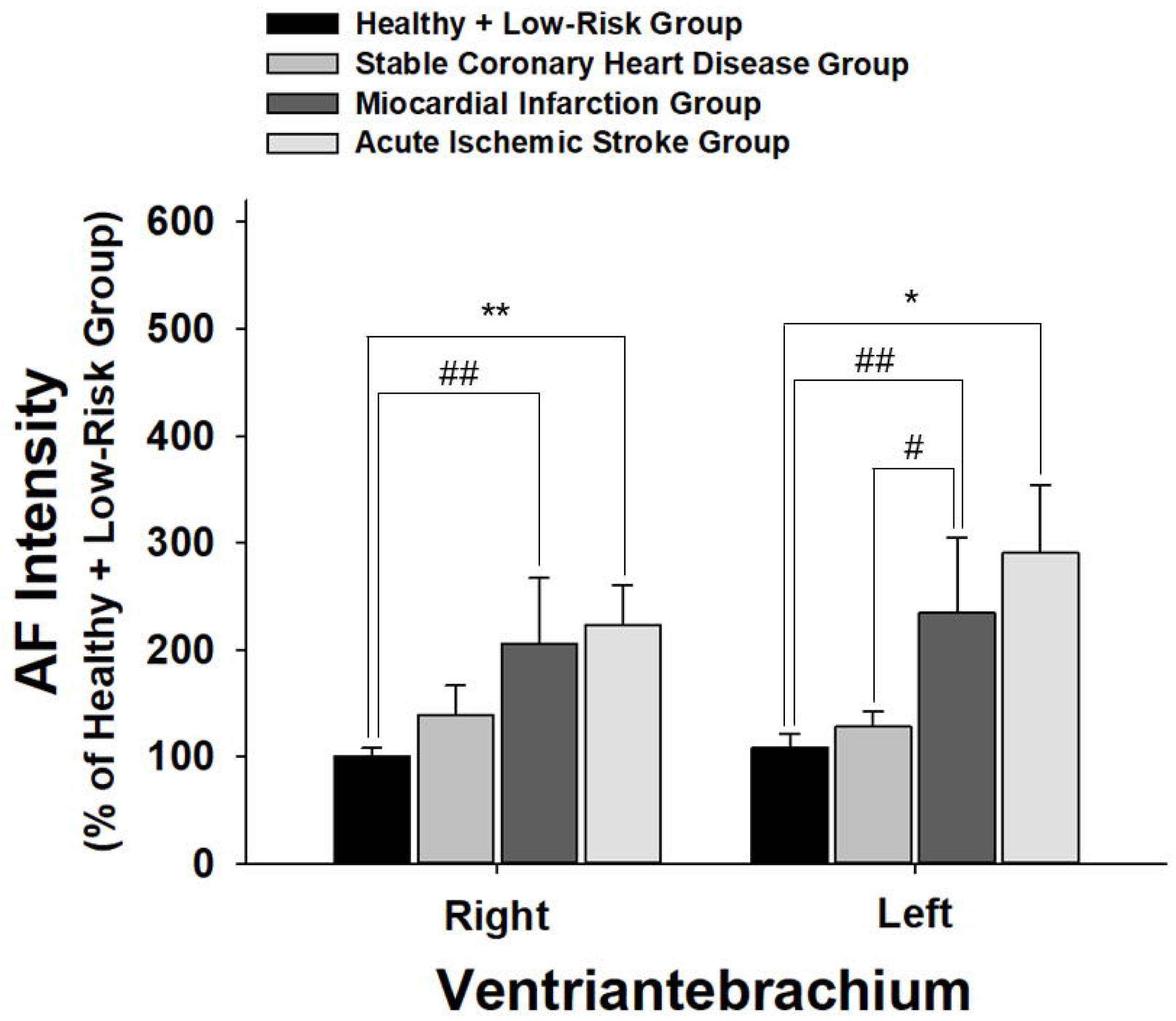
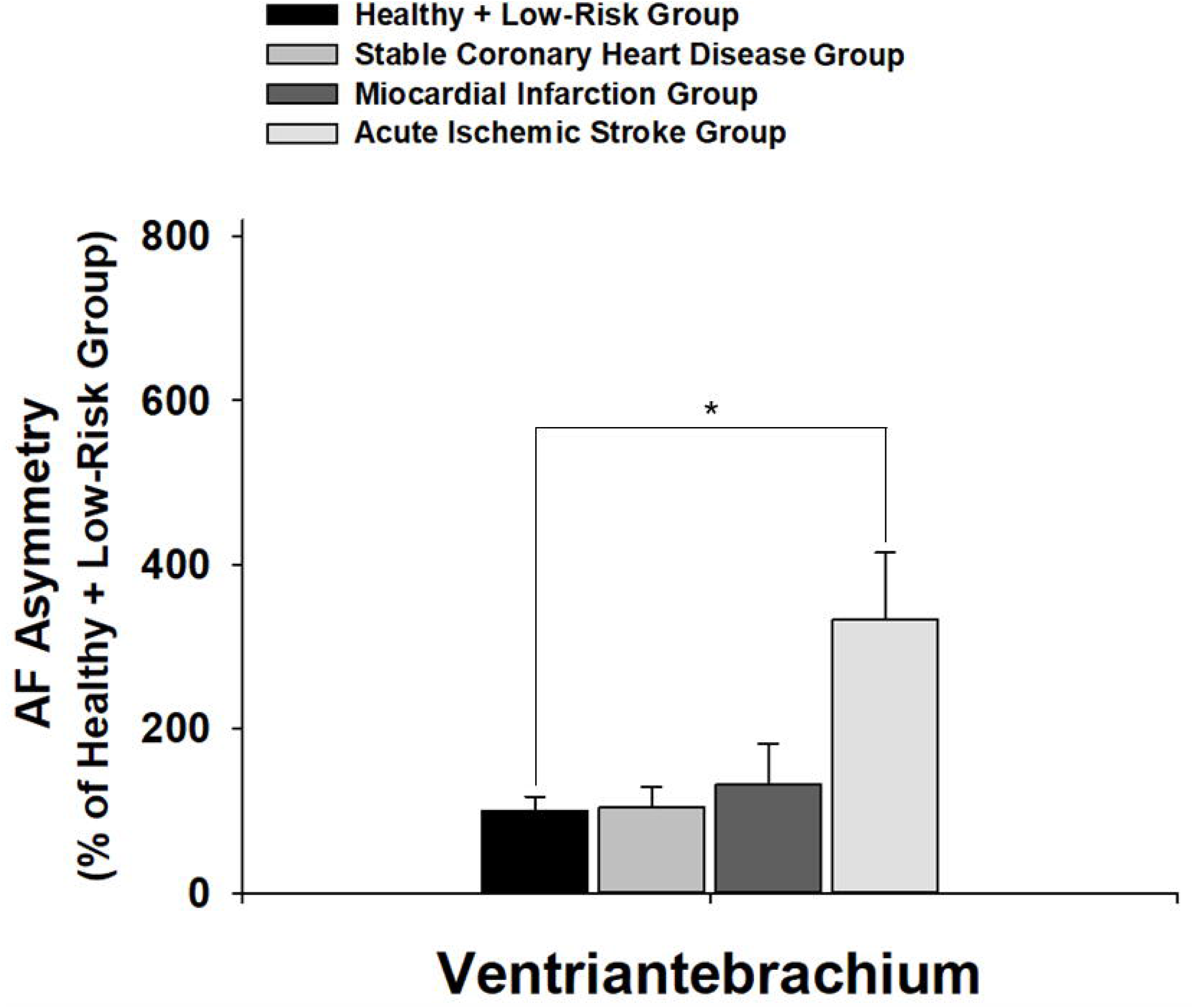
(A) On right and left Ventriantebrachium, both the MI patients and the AIS patients, but not the SCAD patients, have significantly higher green AF intensity than Healthy and Low-Risk persons. (B) Only the AIS patients have significantly higher AF asymmetry than the Healthy and Low-Risk persons at the Ventriantebrachium. The number of subjects in the Healthy and Low-Risk group, the SCAD group, the MI group and the AIS group is 43, 20, 8, and 46-47, respectively. *, *p* < 0.05; **, *p* < 0.01; #, *p* < 0.05 (Student *t-t*est); ##, *p* < 0.01 (Student *t*-test).

In six out of the fourteen regions examined, the AF intensity of the SCAD patients is not significantly different from that of the Healthy and Low-Risk group (Table 1), while in three out of the fourteen regions examined, the AF intensity of the MI patients is not significantly different from that of the Healthy and Low-Risk persons (Table 1). In contrast, only in one out of the fourteen regions examined, the AF intensity of the AIS patients is not significantly different from that of the Healthy and Low-Risk persons (Table 1).

**Table 1.**
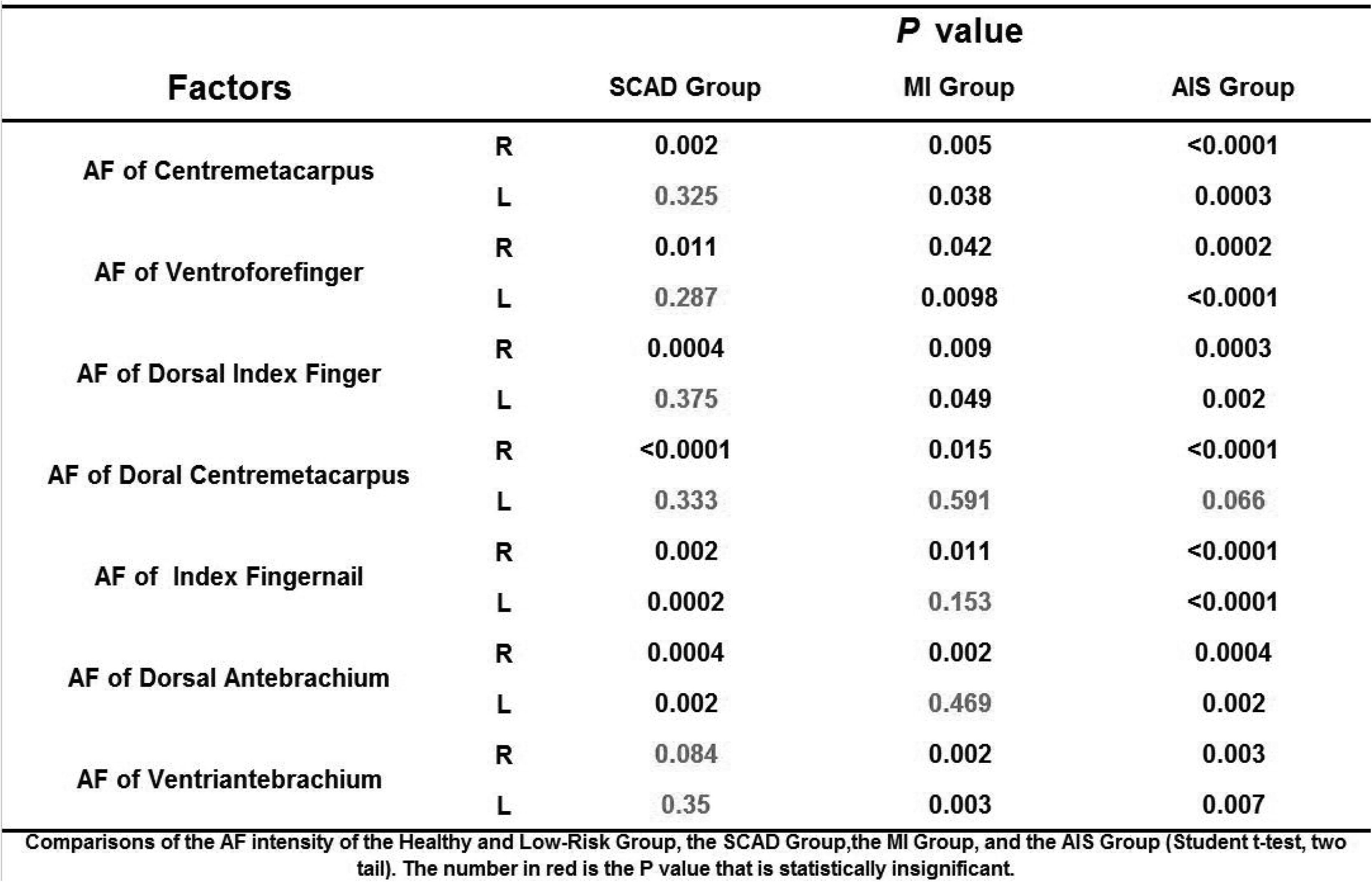
In six out of the fourteen regions examined, the AF intensity of SCAD patients is not significantly different from that of the Healthy and Low-Risk group, while in three out of the fourteen regions examined, the AF intensity of AMI patients is not significantly different from that of the Healthy and Low-Risk group. In contrast, only in one out of the fourteen regions examined, the AF intensity of acute ischemic stroke patients is not significantly different from that of the Healthy and Low-Risk group. The number of subjects in the Healthy and Low-Risk group, the SCAD group, the MI group and the AIS group is 43, 20, 8, and 46-47, respectively.

In three out of the seven positions examined in the left and right side of the body, neither the AF asymmetry of the SCAD group nor the AF asymmetry of the MI group is significantly different from that of the Healthy and Low-Risk group (Table 2). In contrast, in all of the seven positions examined in the left and right side of the body, the AF asymmetry of the AIS group is significantly different from that of the Healthy and Low-Risk group (Table 2).

**Table 2.** In three out of the seven positions examined in the left and right side of the body, neither the AF of the SCAD patients nor the AF of the AMI patients are significantly different from that of the Healthy and Low-Risk group. In contrast, in all of the seven positions examined in the left and right side of the body, the AF asymmetry is significantly different from that of the Healthy and Low-Risk group. The number of subjects in the Healthy and Low-Risk group, the SCAD group, the MI group and the AIS group is 43, 20, 8, and 46-47, respectively.

| Factors | *P* value |  |  |
| --- | --- | --- | --- |
|  | SCAD Group | MI Group | AIS Group |
| Asymmetry of Centremetacarpus | 0.111 | 0.031 | 0.0002 |
| Asymmetry of Ventroforefinger | <0.0001 | 0.0002 | <0.0001 |
| Asymmetry of Dorsal Index Finger | 0.009 | 0.052 | 0.004 |
| Asymmetry of Dorsal Centremetacarpus | 0.009 | 0.097 | 0.003 |
| Asymmetry of Index Fingernail | 0.021 | 0.035 | <0.0001 |
| Asymmetry of Dorsal Antebrachium | 0.051 | 0.004 | 0.0009 |
| Asymmetry of Ventriantebrachium | 0.903 | 0.486 | 0.009 |

Comparisons of the AF asymmetry of the Healthy and Low-Risk Group, the SCAD Group,the MI Group, and the AIS Group (Student t-test, two tail). The number in red is the *P* value that is statistically insignificant.

## Discussion

Our current study has indicated that the MI patients and the SCAD patients have distinct patterns of AF changes, compared with that of the Healthy and Low-Risk persons: First, the AF intensity of the MI patients is significantly higher than that of the Healthy and Low-Risk group in their right and left Centremetacarpus, Ventroforefinger, Dorsal Index Finger and Ventribrachium, while the AF intensity of the SCAD patients is significantly higher than that of the Healthy and Low-Risk group in their right and left fingernails and Dorsal Antebrachium. Second, the AF asymmetry of the MI patients is significantly higher than that of the Healthy and Low-Risk group in their Centremetacarpus, Ventroforefinger, Index Fingernails and Dorsal Antebrachium, while the AF asymmetry of the SCAD patients is significantly higher than that of the Healthy and Low-Risk group in their Ventroforefinger, Dorsal Index Finger, Dorsal Centremetacarpus and Index Fingernails.

Our study has also indicated that AIS patients have markedly differential AF patterns, compared with those of SCAD and MI patients: The percentages of the regions with increased AF intensity of the AIS, MI and SCAD patients are in the following order: AIS patients > MI patients > SCAD patients. Moreover, the percentages of the positions with AF asymmetry of the AIS, MI and SCAD patients are in the following order: AIS patients > MI patients = SCAD patients.

Collectively, these findings have suggested that the distinct patterns of their AF intensity and AF asymmetry could become novel diagnostic biomarkers for MI, SCAD and AIS. These patterns could be the unique ‘Fingerprints of the body surface’s AF’ of the patients. Based on these findings, we proposed a new criteria for MI and SCAD diagnosis – ‘Distinct AF patterns of the body’ Surface’, which may be used jointly with the clinical symptom-based diagnostic criteria for MI and SCAD diagnosis.

Compared with the current diagnostic approaches for MI, our AF-based approach has unique merits: It is non-invasive and economic, and it could be conducted at homes with the applications of big data technology and AI technology. It is expected that the specificity and sensitivity of this diagnostic approach would be further increased, with future increases in the regions examined as well as applications of AI and big data technology. Future studies are needed to further investigate the characteristic merits of our AF-based diagnostic approach compared with current diagnostic approaches for MI.

The oxidative stress induced by UVC has been shown to mediate the increase in the epidermal green AF of mouse ears by inducing keratin 1 proteolysis (7). Because there are significant increases in oxidative stress in the plasma, serum or urine of SCAD patients (1,2,5,6,20) and MI patients (3,4,9,13,20), we propose that the increased oxidative stress may induce increases in the epidermal AF of the SCAD and the MI patients by inducing keratin 1 proteolysis. It is warranted to further test the validity of this proposal.

Our current study has found that the percentage of the regions with increased AF intensity of the MI patients is markedly higher than the SCAD patients. This finding is consistent with our proposal that the oxidative stress mediates the AF increases, since MI patients have significantly higher levels of oxidative stress in their blood than SCAD patients (20). It has also been suggested that plasma concentrations of myeloperoxidase may be used to predict mortality after MI (13). Since the AF increases may result from the oxidative stress in the plasma of MI patients, we propose that our AF-based diagnosis might also be used to predict mortality after MI. Future studies are warranted to determine the validity of this proposal.

## Acknowledgment

The authors would like to acknowledge the financial support by Major Research Grants from the Scientific Committee of Shanghai Municipality #16JC1400500 and #16JC1400502 (to W.Y.) and Chinese National Natural Science Foundation Grant #81271305 (to W. Y.).

## References

1. Baseri M, Heidari R, Mahaki B, Hajizadeh Y, Momenizadeh A, Sadeghi M. Myeloperoxidase levels predicts angiographic severity of coronary artery disease in patients with chronic stable angina. Adv Biomed Res 3: 139, 2014.

2. Bilinska M, Wolszakiewicz J, Duda M, Janas J, Beresewicz A, Piotrowicz R. Antioxidative activity of sulodexide, a glycosaminoglycan, in patients with stable coronary artery disease: a pilot study. Med Sci Monit 15: CR618–23, 2009.

3. Caimi G, Canino B, Incalcaterra E, Ferrera E, Montana M, Lo Presti R. Behaviour of protein carbonyl groups in juvenile myocardial infarction. Clin Hemorheol Microcirc 53: 297–302, 2013.

4. Deepa M, Pasupathi P, Sankar KB, Rani P, Kumar SP. Free radicals and antioxidant status in acute myocardial infarction patients with and without diabetes mellitus. Bangladesh Med Res Counc Bull 35: 95–100, 2009.

5. Kamezaki F, Yamashita K, Tasaki H, Kume N, Mitsuoka H, Kita T, Adachi T, Otsuji Y. Serum soluble lectin-like oxidized low-density lipoprotein receptor-1 correlates with oxidative stress markers in stable coronary artery disease. Int J Cardiol 134: 285–7, 2009.

6. Kiliszek M, Maczewski M, Styczynski G, Duda M, Opolski G, Beresewicz A. Low-density lipoprotein reduction by sirrivastatin is accompanied by angiotensin II type 1 receptor downregulation, reduced oxidative stress, and improved endothelial function in patients with stable coronary artery disease. Coronary Artery Disease 18: 201–209, 2007.

7. Zhang DTM M, H He, Y Li, W Yan, W Yan, Y Zhu, W Ying. UV-Induced Keratin 1 Proteolysis Mediates UV-Induced Skin Damage. bioRxiv, 2017.

8. Maddox TM, Stanislawski MA, Grunwald GK, Bradley SM, Ho PM, Tsai TT, Patel MR, Sandhu A, Valle J, Magid DJ, Leon B, Bhatt DL, Fihn SD, Rumsfeld JS. Nonobstructive coronary artery disease and risk of myocardial infarction. JAMA 312: 1754–63, 2014.

9. Madole MB, Bachewar NP, Aiyar CM. Study of oxidants and antioxidants in patients of acute myocardial infarction. Adv Biomed Res 4: 241, 2015.

10. Mathers CD, Loncar D. Projections of global mortality and burden of disease from 2002 to 2030. PLoS Med 3: e442, 2006.

11. Meerwaldt R, Links T, Graaff R, Thorpe SR, Baynes JW, Hartog J, Gans R, Smit A. Simple noninvasive measurement of skin autofluorescence. Ann N Y Acad Sci 1043: 290–8, 2005.

12. Mingchao Zhang DTM, Yujia Li, Weihai Ying. Oxidative Stress Mediates UVC-Induced Increases in Epidermal Autofluorescence of C57 Mouse Ears. BioRxiv, 2018.

13. Mocatta TJ, Pilbrow AP, Cameron VA, Senthilmohan R, Frampton CM, Richards AM, Winterbourn CC. Plasma concentrations of myeloperoxidase predict mortality after myocardial infarction. J Am Coll Cardiol 49: 1993–2000, 2007.

14. Moran C, Munch G, Forbes JM, Beare R, Blizzard L, Venn AJ, Phan TG, Chen J, Srikanth V. Type 2 diabetes, skin autofluorescence, and brain atrophy. Diabetes 64: 279–83, 2015.

15. O’Gara PT, Kushner FG, Ascheim DD, et al. 2013 ACCF/AHA Guideline for the Management of ST-Elevation Myocardial Infarction A Report of the American College of Cardiology Foundation/American Heart Association Task Force on Practice Guidelines. Journal of the American College of Cardiology 61: E78–E140, 2013.

16. Reichlin T, Hochholzer W, Bassetti S, et al. Early Diagnosis of Myocardial Infarction with Sensitive Cardiac Troponin Assays. New England Journal of Medicine 361: 858–867, 2009.

17. Steg PG, James SK, Atar D, et al., Members ATF. ESC Guidelines for the management of acute myocardial infarction in patients presenting with ST-segment elevation. European Heart Journal 33: 2569–2619, 2012.

18. Thygesen K, Alpert JS, Jaffe AS et al., Task JEAAW. Third Universal Definition of Myocardial Infarction. Circulation 126: 2020, 2012.

19. Thygesen K, Alpert JS, White HD, Force EAAWT. Universal definition of myocardial infarction. Circulation 116: 2634–2653, 2007.

20. Uppal N, Uppal V, Uppal P. Progression of Coronary Artery Disease (CAD) from Stable Angina (SA) Towards Myocardial Infarction (MI): Role of Oxidative Stress. J Clin Diagn Res 8: 40–3, 2014.

21. Vos T, Barber RM, Bell B, et al., Study GBD. Global, regional, and national incidence, prevalence, and years lived with disability for 301 acute and chronic diseases and injuries in 188 countries, 1990-2013: a systematic analysis for the Global Burden of Disease Study 2013. Lancet 386: 743–800, 2015.

